# Inhibition of SARS-CoV-2 Infection in Human Airway Epithelium with a Xeno-Nucleic Acid Aptamer

**DOI:** 10.1101/2023.09.27.559799

**Authors:** Niayesh Razi, Weizhong Li, Maxinne A. Ignacio, Jeffrey M. Loube, Eva L. Agostino, Xiaoping Zhu, Margaret A. Scull, Jeffrey J. DeStefano

## Abstract

**Background:** SARS-CoV-2, the agent responsible for the COVID-19 pandemic, enters cells through viral spike glycoprotein binding to the cellular receptor, angiotensin-converting enzyme 2 (ACE2). Given the lack of effective antivirals targeting SARS-CoV-2, we previously utilized systematic evolution of ligands by exponential enrichment (SELEX) and selected fluoro-arabino nucleic acid (FANA) aptamer R8-9 that was able to block the interaction between the viral receptor-binding domain and ACE2.

**Methods:** Here, we further assessed FANA-R8-9 as an entry inhibitor in contexts that recapitulate infection *in vivo*.

**Results:** We demonstrate that FANA-R8-9 inhibits spike-bearing pseudovirus particle uptake in cell lines. Then, using an *in-vitro* model of human airway epithelium (HAE) and SARS-CoV-2 virus, we show that FANA-R8-9 significantly reduces viral infection when added either at the time of inoculation, or several hours later. These results were specific to the R8-9 sequence, not the xeno-nucleic acid utilized to make the aptamer. Importantly, we also show that FANA-R8-9 is stable in HAE culture secretions and has no overt cytotoxic effects.

**Conclusions:** Together, these results suggest that FANA-R8-9 effectively prevents infection by specific SARS-CoV-2 variants and indicate that aptamer technology could be utilized to target other clinically-relevant viruses in the respiratory mucosa.

## Background

Severe acute respiratory syndrome coronavirus 2 (SARS-CoV-2), the causative agent of COVID-19, has had a devastating impact on society, resulting in nearly 7 million deaths and over 750 million reported infections worldwide. Through vaccination, natural virus evolution, surveillance, and better treatment protocols, the pandemic has dramatically declined, but there are still hundreds of thousands of reported infections each week [1]. Despite noted successes, drug therapies and available vaccines have only served to mitigate COVID-19. This, coupled with the uncertain evolution of the virus, necessitates a continued search for new and improved therapeutics and vaccines that may be applied to SARS-CoV-2 or other coronaviruses that may arise in the future.

A novel approach being pursued by several groups is the development of aptamers that can be employed as therapeutics or in diagnostics. Aptamers are short nucleic acid-based sequences that bind targets with high affinity and are typically produced using systematic evolution of ligands by exponential enrichment (SELEX) [2,3]. Aptamers have the potential to block virus entry into cells or detect specific viruses in diagnostic assays (Reviewed in [4,5]). Several groups have produced DNA aptamers that bind to the SARS-CoV-2 S1 protein (a subunit of the viral spike protein found on the outer surface of the virus envelope) and block the interaction between the receptor binding domain (RBD) of S1 and the host cell ACE2 receptor [6–16; Reviewed in: 17,18]. Blocking this interaction prevents virus entry into cells, similar to how current neutralizing antibody therapies block SARS-CoV-2 infection (Reviewed in [19]). We previously reported the analysis of aptamer FANA-R8-9, which was made from 2’-fluoro-arabino nucleic acid (FANA) and was able to block the interaction between the viral RBD and ACE2 [15]. FANA is a xeno-nucleic acid (XNA), where XNAs are synthetic nucleic acids with altered sugar, base, or phosphate backbones. These alternatives to DNA and RNA aptamers often have advantageous properties as they may accommodate unique structures, have low immunogenicity, and have greater resistance to degradation [19–24].

Still, whether FANA-R8-9 can block spike-ACE2 interaction in the context of a viral particle, and whether FANA-R8-9 – or any spike-targeting aptamer, to our knowledge – can function to block SARS-CoV-2 infection in the native respiratory microenvironment remains unknown. Notably, human airway epithelial (HAE) cultures at air-liquid interface have been established and recapitulate both the morphology and physiology of the airway epithelium *in vivo,* including the establishment of a mucus barrier [25,26]. Importantly, these cultures are susceptible to SARS-CoV-2 and have been previously used to assess the efficacy of other antivirals against SARS-CoV-2 replication *in vitro* [27,28]. Here, we report that FANA-R8-9 not only inhibits the uptake of spike-bearing pseudovirus particles, but also blocks SARS-CoV-2 infection in the HAE system, demonstrating the potential utility of XNA aptamers to target respiratory viruses in the mucosal microenvironment.

## Materials and Methods

### Generation of aptamers

FANA-R8-9 was previously selected for binding to the SARS-CoV-2 spike RBD (Arg319-Phe541; Wuhan strain) through SELEX [15]. FANA-R8-9 and various controls were produced by enzymatic synthesis using DNA templates and D4K polymerase [21], as previously described [15]. The DNA templates were 18 nucleotides longer than the products, allowing separation on denaturing acrylamide gels. Sequences are shown below with underlined bases composed of FANA and non-underlined DNA. FANA-R8-9: 5′-AAAAGGTAGTGCTGAATTCG<u>CGAGCCCGCAUGAAAAGGGGAGAUAAAAAAUAUCUGUCGAUUCGCUAUCCAGUUGGCCU</u>-3’; Scrambled FANA: 5′-AAAAGGTAGTGCTGAATTCG<u>(N)_40_UUCGCUAUCCAGUUGGCCU</u>-3’ (N-FANA A, U, C or G); Fixed FANA (an unrelated sequence derived from the starting pool for SELEX): 5’-AAAAGGTAGTGCTGAATTCG<u>AGUCACGCCAACACAGGAAGCGUAGGUCUAUCUUGUAGGUUUCGCUAUCCAGUUGGCCU</u>-3’; Fixed DNA (A DNA only copy of Fixed FANA): 5’AAAAGGTAGTGCTGAATTCGAGTCACGCCAACACAGGAAGCGTAGGTCTATCTTGTAGGTTTCGCTATCCAGTTGGCCT-3’; IN-1.1 (FANA HIV-1 integrase aptamer [29]: 5’-AAAAGGTAGTGCTGAATTCG<u>UUUCAAGUGUAUAUUAACUACGCAUCUUUCCCCCUGCGUAUUCGCUAUCCAGUUGGCCU</u>-3’. After extracting sequences from gels and recovery, the material was run over a Sephadex G-25 column to remove impurities. All sequences were quantified using absorbance at 260 nM assuming that FANA nucleotides absorb like DNA.

### Cell lines and human airway epithelial cultures

Vero-E6 (with high expression of endogenous ACE2, Cat No. NR-53726) were obtained from Biodefense and Emerging Infections Research Resources Repository (BEI Resources, Manassas, VA). Vero-E6 cells were maintained in complete Dulbecco’s Minimal Essential Medium (DMEM; Invitrogen Life Technologies), supplemented with 10% fetal bovine serum (FBS) only, or 10% FBS plus 2 mM L-glutamine, nonessential amino acids, 100 units/ml of penicillin, 100 μg/ml of streptomycin, and 250 ng/ml of amphotericin B. Vero-E6 cells routinely tested negative for Mycoplasma sp. by real-time PCR.

Human airway tracheobronchial epithelial cells isolated from airway specimens from donors without underlying lung disease were provided by Lonza, Inc. Primary airway cells derived from single patient sources were first expanded on plastic in Pneumacult-Ex Plus medium (Cat No. 05040, StemCell Technologies), then seeded (3.3 × 10^4^ cells / well) on rat tail collagen type 1-coated permeable Transwell® membrane supports (6.5 mm; Cat No. 3470, Corning, Inc.). Cultures were differentiated in Pneumacult-ALI medium (Cat No. 05001, StemCell Technologies) with the provision of an air-liquid interface for approximately 6 weeks to form polarized cultures that resemble *in-vivo* pseudostratified mucociliary epithelium. All cell cultures were maintained at 37°C with 5% CO_2_. All experiments involving HAE cultures utilized at least two different donors, and the data in the figures are indicative of unique biological replicates.

### Generation of spike-bearing pseudovirus particles and SARS-CoV-2 virus

The spike-bearing pseudovirus particles (rVSVΔG/SARS-CoV-2-S-D614Gd21-NLucP; Cat No. EGA298-PM, Kerafast, Inc.) were previously described [30] and obtained commercially for this study. In this recombinant vesicular stomatitis virus, the native glycoprotein gene has been replaced with the SARS-CoV-2 spike gene from the Wuhan strain lacking the last 21 residues of the cytoplasmic tail and containing an amino acid change (D614G) to promote the incorporation of the spike protein. This virus was propagated once in Vero-E6 cells and quantified by endpoint dilution (Tissue Culture Infectious Dose (TCID_50_)) assay in Vero-E6 cells.

SARS-CoV-2 ancestral strain hCoV-19/USA/NY-PV08410/2020 (referred to as wild type and abbreviated as WT, Cat No. NR-53514) and Delta strain hCoV-19/USA/PHC658/2021(B.1.167.2) (Cat No. NR-55611) were obtained from BEI Resources with the permission of the Centers for Disease Control and Prevention (CDC). Viruses from BEI Resources were passaged in Vero-E6 cells cultured in a medium with reduced (1%) FBS. At 72 hrs post-infection, tissue culture supernatants were collected and clarified before being aliquoted and stored at −80°C. The virus stocks were titrated using the TCID_50_ assay. All experiments using live SARS-CoV-2 were performed in an approved Animal Biosafety Level 3+ (ABSL3+) facility at the University of Maryland using appropriate powered air-purifying respirators (PAPR) and protective equipment.

### Pseudovirus particle inhibition assays

Vero-E6 cells were seeded in a 96-well plate at a density of 1.4×10^4^ cells / well and incubated overnight at 37°C. The following day, an inoculum containing pseudovirus particles (stock titer = 3×10^6^ TCID_50_ / ml, diluted 1:64 in final inoculum) and FANA-R8-9 or control aptamer (50 nM, with 2-fold dilutions to 0.78 nM final concentration) was prepared and incubated for 30 min at room temperature. The cell culture medium was then removed, and the inoculum (25 μl) was applied directly to the cells, followed by a 2-hour incubation at 37°C. Serum-free DMEM containing FANA-R8-9 or control aptamer at the matching concentration was then added to reach a final 100 μl volume in each well. Twenty-four hours later, luciferase activity in each well was quantified using the Nano-Glo ® Luciferase Assay System (Cat No. N1120; Promega) according to the manufacturer’s protocol and a Berthold Centro XS^3^ LB 960 plate reader. Relative luciferase units (RLU) were then normalized to “no aptamer” to determine the percent inhibition.

### Pseudovirus aptamer binding assay

Pseudovirus particles (stock titer 3×10^6^ TCID_50_ / ml) were concentrated by ultracentrifugation through a 20% sucrose cushion. Briefly, approximately 2 ml of virus in DMEM supplemented with 10% FBS was mixed with 1.5 ml of phosphate buffered saline (PBS, 1.7 mM KH_2_PO_4_, 5 mM Na_2_HPO_4_, 150 mM NaCl, pH 7.4) and loaded into a centrifuge tube (Beckman Cat. No. 344057, 5 ml capacity). One milliliter of 20% sucrose in 20 mM HEPES pH 7.4, 150 mM NaCl was then underlaid into the centrifuge tube, and PBS was added to the top to ensure proper fill. Centrifugation was done for 2 hrs at 35,000 rpm using a Beckman SW 55 Ti rotor. The supernatant was carefully removed and the pellet, along with a small amount of remaining sucrose solution, was resuspended in 200 μl of PBS. Aliquots were stored at −80°C. The binding assay was performed by mixing concentrated pseudovirus (0, 0.63, 1.25, 2.5, or 5 μl) with radiolabeled aptamer or controls (20 nM final concentration) in a total volume of 15 μl PBS. Reactions also contained 15 pmoles of unlabeled DNA competitor (5′-AAAAGGTAGTGCTGAATTCG<u>(N)_40_TTCGCTATCCAGTTGGCCT</u>-3’ (N-DNA A, T, C, or dGTP). Samples were incubated at room temperature for 20 min, then 50 μg of heparin sulfate (in PBS) was added, and incubations were continued for 40 min. Competitor DNA and the heparin post-binding step were necessary to reduce background binding such that only tightly bound nucleic acids would remain associated with virus material. Samples were filtered through pre-wetted (25 mM Tris-HCL pH 7.5, 10 mM KCl) nitrocellulose filters (Whatman™ Protran™, 25 mm diameter, 0.45 μm pore size) and washed with 5 ml of the per-wetting buffer at a rate of ∼0.5 ml/sec. Filters were quantified in a scintillation counter.

### SARS-CoV-2 inhibition assays in HAE

HAE cultures were washed with PBS twice, for 15 min at 37°C each time, on the apical surface, and provided with fresh Pneumacult-ALI medium immediately prior to inoculation. In experiments where virus was mixed with aptamer before inoculation, aptamers were reconstituted in water at 3 pmol / μl. Aptamer (1 μl; or water (vehicle control)) was then mixed with SARS-CoV-2 (29 μl; wild type stock titer was 8.51 × 10^6^ TCID_50_ / ml; Delta stock titer was 5 × 10^6^ TCID_50_ / ml) and incubated for 30 min at room temperature. After this, 20 μl of the inoculum mixture was added to the apical surface of appropriate cultures before returning them to the 37°C incubator. The viral inoculum was left on the culture for the duration of the experiment. At 24 and 48 hrs post-infection (hpi), 0.5 μl of aptamer (or water) was mixed with 14.5 μl PBS and 10 μl added to the apical surface of corresponding cultures. All cultures (including an uninfected (mock) culture run in parallel) were fixed at 72 hrs post-inoculation in paraformaldehyde for 60 min prior to removal from the ABSL3, in alignment with IBC-approved protocols. An additional set of experiments was performed in which aptamer was added after virus inoculation (at 0, 2, or 6 hpi) and again at 24 and 48 hrs post-infection prior to fixing at 72 hrs, as above.

### *En face* immunofluorescence staining

Fixed HAE cultures were washed twice in PBS prior to permeabilization with 2.5% triton X-100 / PBS for 15 min. HAE cultures were then blocked with 3% bovine serum albumin (BSA) / PBS for 1 hr. Primary antibody, specific for the SARS-CoV-2 nucleoprotein (clone 1C7, Cat No. bsm-41411M, Bioss), was then applied to the apical surface at a 1:500 dilution in 1% BSA / PBS and incubated overnight at 4°C. Following three washes with PBS, a secondary antibody (donkey anti-mouse IgG Alexa Fluor 488, Cat No. A21202, Invitrogen) was diluted 1:500 in 1% BSA / PBS and applied to the apical surface for 1 hr at room temperature. Cultures were then washed three times with PBS and viral antigen was visualized, and images were acquired with either a Zeiss Axio Observer 3 inverted fluorescence microscope or a Zeiss Axio Observer Z1 microscope with motorized stage, both equipped with Zeiss AxioCam 503 mono cameras and Zen imaging software, including the Tiles & Positions module. A Fiji macro was used to analyze whole culture composite images, or 4-5 images from pre-determined regions of the culture per replicate well to determine the percent antigen-positive area. The data were exported to Excel, and the average percent antigen-positive area was calculated per replicate. The resulting area quantification was then visualized in GraphPad Prism.

### Aptamer degradation assays

Apical secretions from HAE cultures were harvested by adding 100 μl of PBS to the culture surface and incubating at 37°C for 15 min before collection in a microcentrifuge tube with a pipette. Washes were stored at 4°C before use or used immediately for aptamer degradation testing. Apical washes from several donors were used in independent experiments and yielded similar results. Assays were conducted by adding 5’-^32^P-labeled FANA-R8-9 or DNA-R8-9 (100 nM final concentration) directly to apical wash material in a total volume of 100 μl. A 10 μl aliquot was removed for digestion with DNaseI (2 U) for 2 hrs at 37°C. After digestion, 10 μl of 2X proteinase K digestion buffer (100 mM Tris-HCl pH 7.5, 100 mM NaCl, 20 mM EDTA pH 8, 10 mM DTT, 2% (w:v) SDS) was added to the sample, which was then frozen at −80°C. A second 10 μl was removed as a “time 0” control and mixed with the same buffer. The remaining 80 μl was incubated at 37°C and 10 μl aliquots were removed and placed in buffer at 0.5, 1, 2, 4, 8, or 24 hrs. The sample tube was briefly centrifuged before aliquot removal to bring all liquid to the bottom. Frozen samples were thawed and 20 μg of proteinase K was added and samples were incubated in a PCR machine at 50°C for 8 hours. Twenty μl of 2X gel loading buffer (90% formamide, 10 mM EDTA pH 8, 0.025% each bromophenol blue and xylene cyanol) was added to each sample and samples were electrophoresed on a 10% polyacrylamide-7M urea gel in Tris-Borate-EDTA buffer as described [31]. Gels were dried and then imaged using an Amersham Typhoon phosphorimager.

### Evaluation of cytotoxicity

To assess the direct effect of FANA-R8-9 on HAE culture integrity, cultures were washed twice with PBS and aptamer applied at 0, 24, and 48 hrs, as above (see “SARS-CoV-2 inhibition assays in HAE”). At 72 hrs post-initial aptamer application, 100 μl of PBS was added to the apical surface, incubated for 30 min at 37°C, and then collected and used to quantify lactate dehydrogenase (LDH) release (in a 50 μl aliquot) using the CytoTox 96 Non-Radioactive Cytotoxicity Assay kit (Cat No. G1780, Promega) according to the manufacturer’s protocol. The basolateral media was then replaced with 500 μl PBS and another 200 μl PBS applied to the apical surface before measuring the transepithelial electrical resistance (TEER) using the Millicell Electrical Resistance System (ERS)-2 (Sigma). Following TEER measurements, PBS was removed and HAE cultures were fixed in 4% paraformaldehyde prior to paraffin embedding and sectioning at the New York University Experimental Pathology Research Laboratory (New York, NY). Five micrometer-thick sections on slides were stained with hematoxylin and eosin (H&E) to allow histological evaluation of the epithelium by light microscopy. Images of H&E-stained sections were acquired using a Zeiss Axio Observer Z1 microscope and AxioCam Erc5s camera.

### Statistical analysis

Statistical analysis of resulting data was done using Prism GraphPad v.9 software (GraphPad) and is further detailed in the figure legends.

## Results

### FANA Aptamer R8-9 inhibits SARS-CoV-2 pseudovirus particle uptake in cell lines

We previously demonstrated that FANA aptamer R8-9 could block the binding of ACE2 to the spike protein receptor binding domain [15]. Thus, to further explore the ability of FANA-R8-9 to inhibit SARS-CoV-2 infection, we sought to determine whether FANA-R8-9 could block spike protein-mediated uptake of pseudoviral particles into cells. To do so, we utilized luciferase-expressing, VSV-based pseudovirus particles bearing the spike protein from the Wuhan strain. Pseudovirus particles were mixed with FANA-R8-9, or other aptamer controls, prior to inoculation of Vero-E6 cells, and luciferase expression was measured at 24 hrs post-inoculation as a proxy of successful pseudovirus particle entry. Pseudovirus signal decreased with increasing concentrations of FANA-R8-9, with a near complete block of infection at 50 nM (**Figure 1A**). Various control aptamers also inhibited the pseudovirus signal, with the FANA-IN-1.1 and Fixed FANA aptamers demonstrating higher levels of inhibition than the Fixed DNA aptamer. Since this assay measures the expression of pseudovirus genomic products, both entry and virus replication are required for a signal. Thus, FANA could possibly affect either or both of these steps in the viral life cycle, leading to the inhibition observed with FANA-IN-1.1 and Fixed FANA controls. A direct block of entry is most easily explained by the binding of the aptamer to the pseudovirus spike protein. To test the binding of FANA-R8-9 and control aptamers to the pseudovirus particles, the assay depicted in **Figure 1B** was used. Concentrated pseudovirus particles were mixed and incubated with radiolabeled aptamer along with excess nonspecific DNA (unlabeled) of the same size. Heparin was then added and incubation continued for 40 min before processing the material for binding to nitrocellulose filters, which would retain the labeled aptamer only if bound to protein. The inclusion of nonspecific DNA and incubation with heparin, a binding competitor for nucleic acid, was required to decrease background binding to material in the pseudovirus preparation. Under these conditions, only nucleic acids that bind with high stability to the protein material in the pseudovirus preparation would be measurable. Results showed that FANA-R8-9 bound in a concentration-dependent manner to pseudovirus material while the controls showed weak binding **(Figure 1C**). These data suggest that only FANA-R8-9 interacts strongly with spike protein in the context of a viral particle. The inhibition observed with FANA controls is likely the result of a mechanism that does not involve binding to the pseudovirus spike protein.

**Figure 1.**
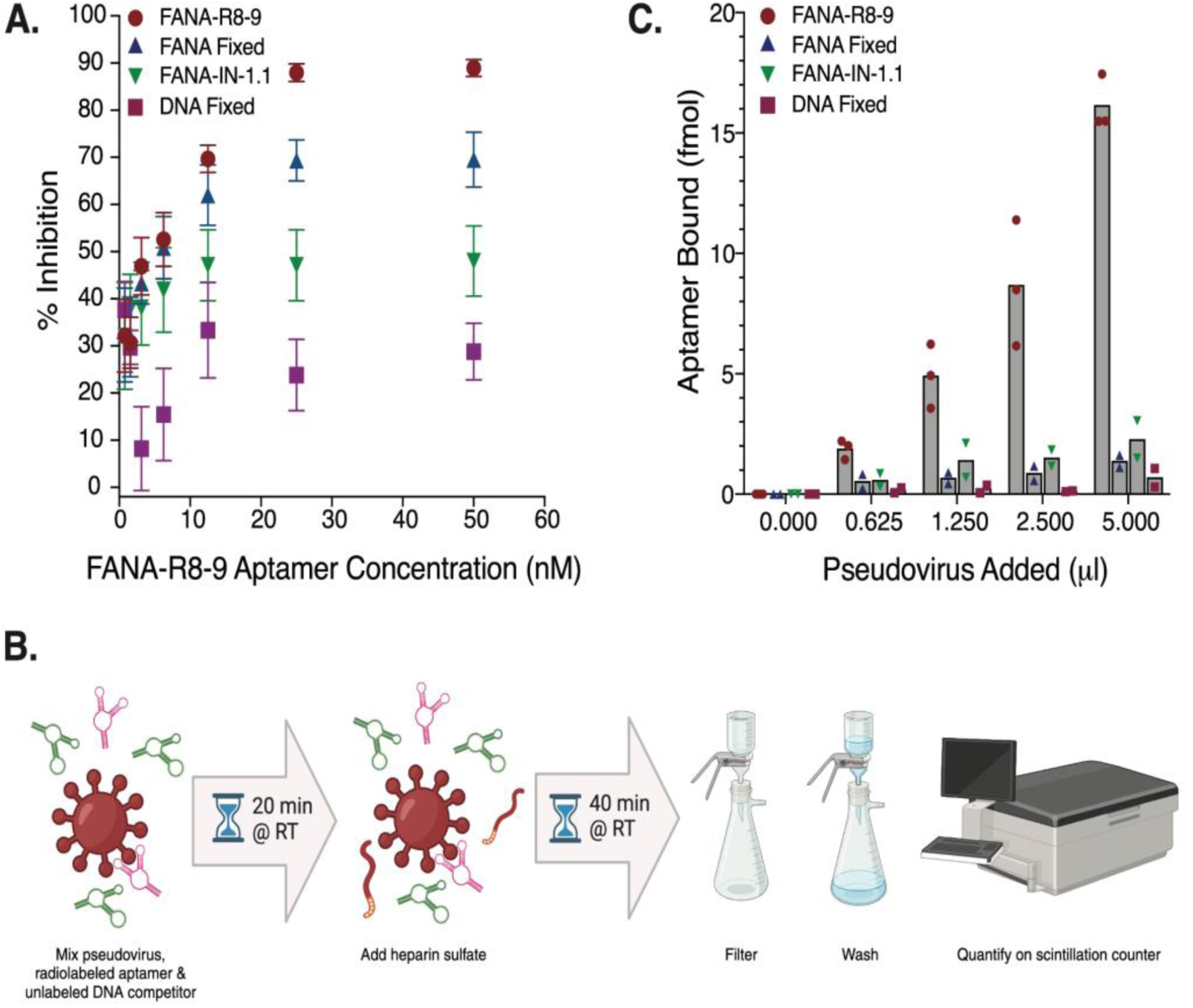
Interaction between FANA-R8-9 and SARS-CoV-2 spike-bearing pseudovirus particles. **A**) Aptamer-mediated inhibition of pseudovirus particle uptake in Vero E6 cells. Data shown represent the mean of 6-12 replicates per condition, combined across three (Fixed DNA; FANA-IN-1.1), four (Fixed FANA), or six (FANA-R8-9) experiments +/− SEM. **B**) Cartoon schematic of the experimental workflow utilized to generate the data in panel C. Radiolabeled FANA-R8-9 (pink); unlabeled DNA competitor (green). RT = room temperature. Created with BioRender.com **C**) Quantification of FANA-R8-9 (or control) aptamer bound to increasing amounts of SARS-CoV-2 spike-bearing pseudovirus particles. Data shown represent the mean of single replicates per condition, combined across two (Fixed FANA; FANA-IN-1.1; Fixed DNA) or three (FANA-R8-9) experiments.

### FANA-R8-9 inhibits SARS-CoV-2 infection in HAE cultures

Given the ability of FANA-R8-9 to bind pseudovirus particles harboring the SARS-CoV-2 spike protein and inhibit their entry, we next asked if this aptamer could block SARS-CoV-2 virus infection in a model of human airway epithelium that recapitulates infection in a more physiologically-relevant setting. Normal human tracheobronchial cells from individual donors were differentiated on Transwell® membrane supports at air-liquid interface and mucus secretions were removed prior to inoculation to facilitate infection. Preliminary tests indicated that FANA-R8-9 concentrations less than 100 nM were associated with stochastic SARS-CoV-2 inhibition in HAE (data not shown); thus, 100 nM aptamer was utilized in all subsequent assays. In initial experiments, 100 nM FANA-R8-9, or water as a vehicle control, was mixed with virus prior to inoculation of the apical culture surface (with an approximate multiplicity of infection (MOI) of 1). Additional aptamer was added to corresponding cultures at 24 and 48 hpi, and the extent of infection across the epithelium was evaluated at 72 hpi following *en face* staining for viral antigen and fluorescence microscopy (**Figure 2A**). To further assess the ability of the aptamer to inhibit more recent variants, we performed these experiments with both the wild-type strain (hCoV-19/USA/NY-PV08410/2020) and Delta variant (hCoV-19/USA/PHC658/2021(B.1.167.2)). Notably, FANA-R8-9 significantly reduced the nucleocapsid-positive area in HAE following infection with either strain, indicating fewer cells were infected in the presence of the aptamer (**Figure 2B, D**).

**Figure 2.**
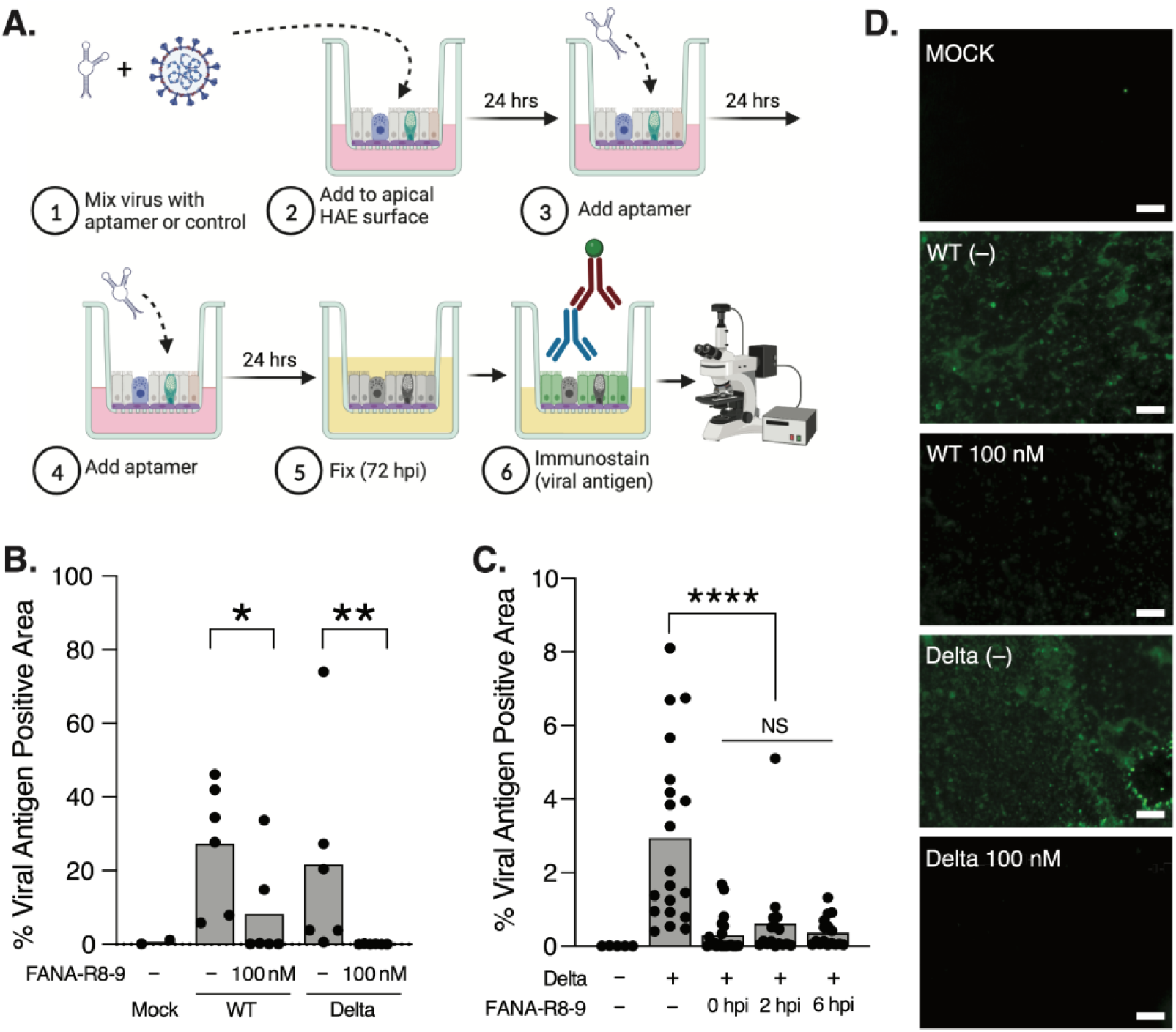
FANA-R8-9 aptamer inhibits SARS-CoV-2 infection in HAE cultures. **A**) Cartoon schematic of the experimental workflow used to obtain the data in panel B. Created with BioRender.com. Hours post-infection = hpi. **B**) The percent viral antigen positive area, quantified following *en face* immunofluorescence staining and imaging in HAE cultures infected with WT SARS-CoV-2 or Delta variant in the presence or absence of FANA-R8-9 aptamer. Individual biological replicates across two experiments (n=3 per donor; 2 donors) are shown with the bar representing the mean. Experimental results were analyzed by the Mann-Whitney test between indicated conditions. All data are significant where indicated (*p < 0.05; ** p<0.005). **C**) The percent viral antigen positive area determined as in panel B for cultures that were first inoculated with the Delta variant, then subsequently treated with vehicle or FANA-R8-9 aptamer immediately following virus inoculation (0), or 2, or 6 hpi. FANA-R8-9 aptamer was added to the indicated cultures again at 24 and 48 hpi prior to fixation at 72 hpi and analysis by *en face* immunostaining and fluorescence microscopy as in panels A and B. Individual biological replicates for one experiment (n=4 per donor; 4-5 donors per condition) are shown with the bar representing the mean. Experimental results were analyzed using ordinary one-way ANOVA between indicated conditions. Data are non-significant (NS) or significant where indicated (**** p < 0.0001). **D**) Representative photomicrographs of HAE cultures analyzed in panel B. HAE were fixed at 72 hpi and immunoprobed *en face* for the presence of SARS-CoV-2 nucleoprotein (green) after apical inoculation with vehicle (mock) or SARS-CoV-2 (WT or Delta variant) in the presence (100 nM) or absence (–) of FANA-R8-9 aptamer. Scale bar = 100 microns.

We next sought to assess the ability of FANA-R8-9 to act in a “therapeutic” setting, by adding the aptamer directly to the apical surface of HAE cultures after virus inoculation. We selected the Delta variant for these assays since FANA-R8-9 was able to completely block infection with this virus in our initial experiments (**Figure 2B**). Delta was added to the apical surface of HAE and FANA-R8-9 was applied either immediately (0), or 2, or 6 hpi, followed by additional aptamer treatments at 24 and 48 hpi. While infection could be detected at 72 hpi in some FANA-R8-9-treated cultures, the viral nucleoprotein-positive area in FANA-R8-9-treated HAE was still significantly reduced compared to HAE without aptamer treatment (**Figure 2C**). These data indicate that FANA-R8-9 can interact with SARS-CoV-2 and limit infection within the airway mucosal environment even several hours after viral infection. The treatment window was not indefinite, however, since adding FANA-R8-9 to Delta-inoculated HAE cultures 24 hpi did not significantly impact the infection (data not shown).

### Inhibition of SARS-CoV-2 infection in HAE is specific to FANA-R8-9

In our pseudovirus particle inhibition assays (**Figure 1A**), we noted that other aptamers could limit pseudovirus particle uptake and / or luciferase expression in Vero-E6 cells, though not to the same extent as FANA-R8-9. To verify the specificity of Delta variant inhibition by FANA-R8-9 in HAE, we assayed FANA fixed, FANA scramble, and DNA fixed aptamers in parallel with FANA-R8-9 in the Figure 2A workflow. Consistent with our previous results (**Figure 2B, 2C**), FANA-R8-9 completely blocked Delta variant infection (**Figure 3**) when the viral antigen-positive area was assessed at 72 hpi. In contrast, none of the other control aptamers significantly affected SARS-CoV-2 infection indicating that FANA-R8-9-mediated inhibition is specific to the FANA-R8-9 sequence.

**Figure 3.**
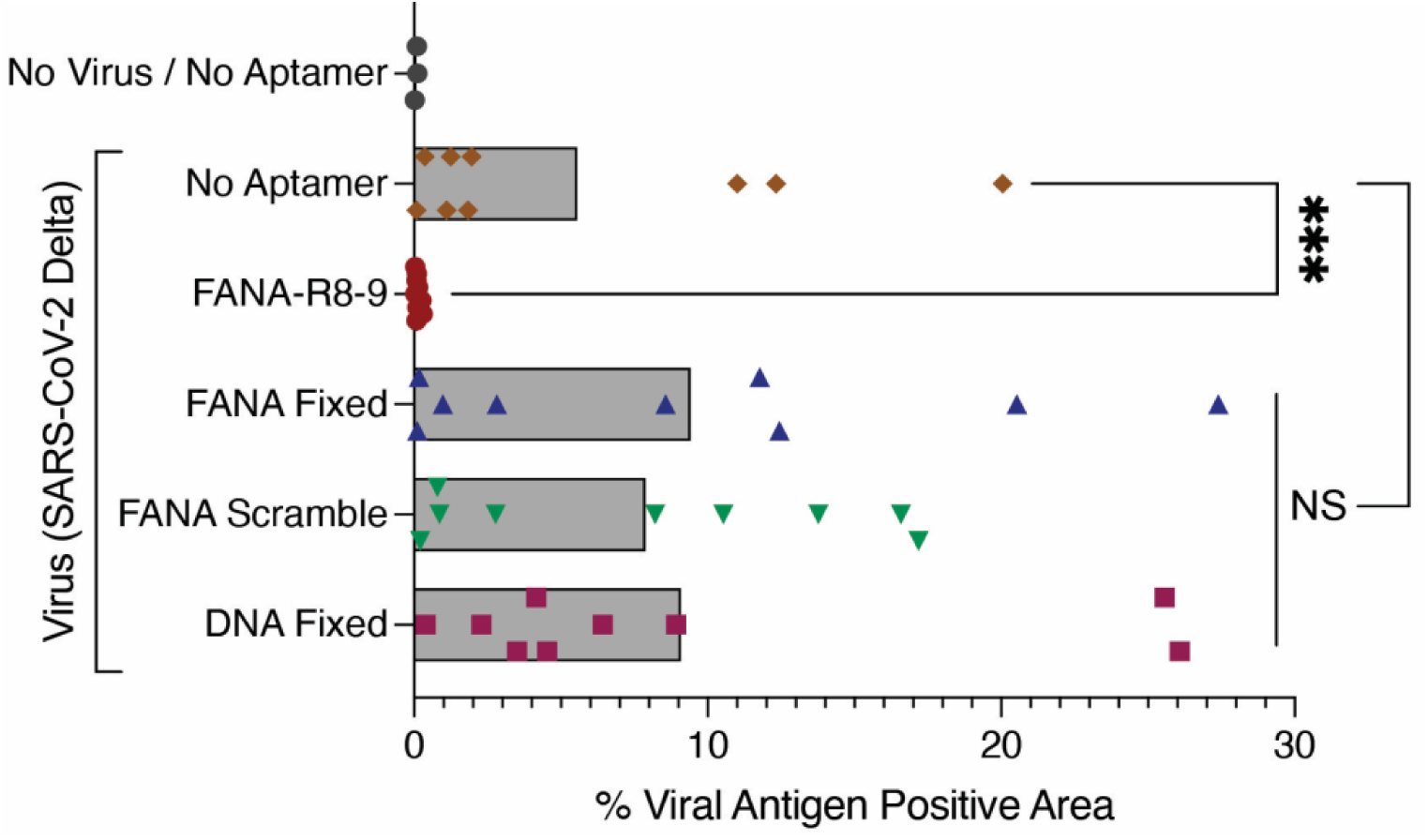
FANA-R8-9 aptamer-mediated inhibition of SARS-CoV-2 infection in HAE cultures is sequence-specific. HAE cultures were inoculated with the indicated treatments at 0, 24, and 48 hrs, (as outlined in Figure 2A), then fixed at 72 hrs. The percentage of viral antigen-positive area was then quantified following *en face* immunofluorescence staining and imaging of SARS-CoV-2 nucleoprotein in fixed cultures. Individual biological replicates are shown (n=3 replicates per condition, per donor; 2 donors (1 donor was assayed twice, in independent experiments) with the bar representing the mean. Experimental results were analyzed using the Mann-Whitney test (between No Aptamer and FANA-R8-9 conditions) or one-way ANOVA (between No Aptamer and all aptamer controls) and data are non-significant (NS) or significant (*** p < 0.001) where indicated.

### FANA-R8-9 is stable in apical secretions from HAE cultures and does not induce overt toxicity

We previously demonstrated that FANA-R8-9 remained intact for several hrs in serum-containing cell culture media, and had a breakdown rate approximately equal to a similar DNA aptamer [15]. To assess FANA-R8-9 stability in airway mucus, we harvested apical secretions from HAE cultures and monitored FANA-R8-9 decay over 24 hrs in these secretions, or in DMEM culture media supplemented with 10% FBS for comparison (**Figure 4**). The latter condition mimics the previous results noted above and demonstrates modest, though clear decay over 24 hrs. In contrast, FANA-R8-9 was more stable in HAE secretions, showing a very slight breakdown only after 24 hrs. The results indicate that FANA-R8-9 retains potency in HAE cultures from several hrs to days.

**Figure 4.**
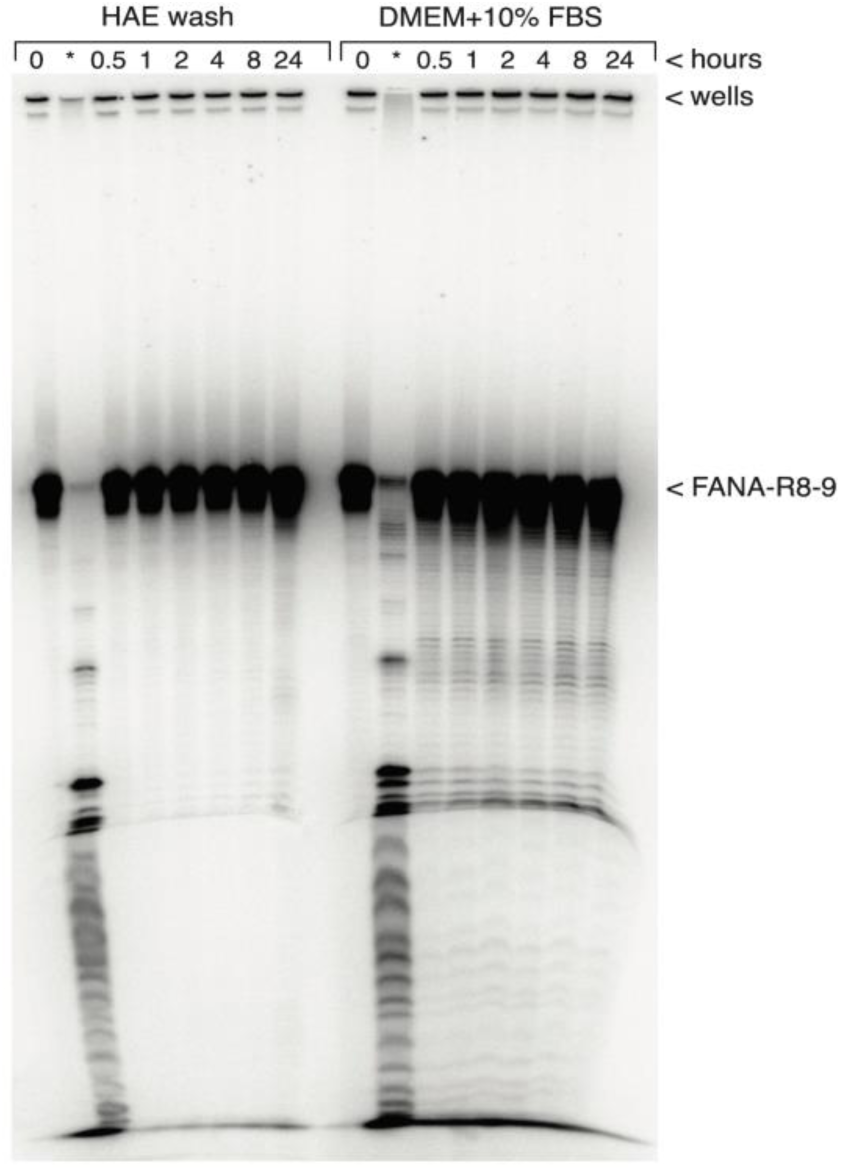
FANA-R8-9 aptamer is stable in apical secretions from HAE cultures. Degradation of FANA-R8-9 aptamer in HAE and serum-supplemented DMEM. Radiolabeled FANA-R8-9 aptamer was incubated in HAE washes or serum-supplemented DMEM medium as described in Materials and Methods. Aliquots were removed at the indicated time points and processed as described, then run on a 10% denaturing polyacrylamide-urea gel. The dried gel was exposed and imaged with a phosphorimager. A small amount of material remained in the wells during electrophoresis. The position of the aptamer (∼79 nts) is indicated. * An aliquot of the sample was digested with 2 U of DNase I for 2 hrs in the indicated medium.

Given the lack of FANA-R8-9 degradation in HAE secretions, we next sought to determine if FANA-R8-9 demonstrated any overt toxicity in HAE cultures using two types of cell toxicity assays, conducted with 100 and 1000 nM aptamer. The latter concentration is 10X greater than the maximum concentration in virus inhibition assays. HAE cultures were first treated with aptamer using the same timing and conditions employed in infection assays. This was followed by toxicity assays that use lactate dehydrogenase (LDH) release as a proxy for cell death and transepithelial electrical resistance (TEER) as a measure of epithelial barrier integrity. Notably, both LDH release (**Figure 5A**) and TEER (**Figure 5B**) assays demonstrated no significant toxicity at 100 or 1000 nM aptamer. These results were further supported by a lack of overt cytopathology in histological cross-sections of HAE and are consistent with the low toxicity observed with FANA oligonucleotides in cell culture by others (**Figure 5C**) [32,33].

**Figure 5.**
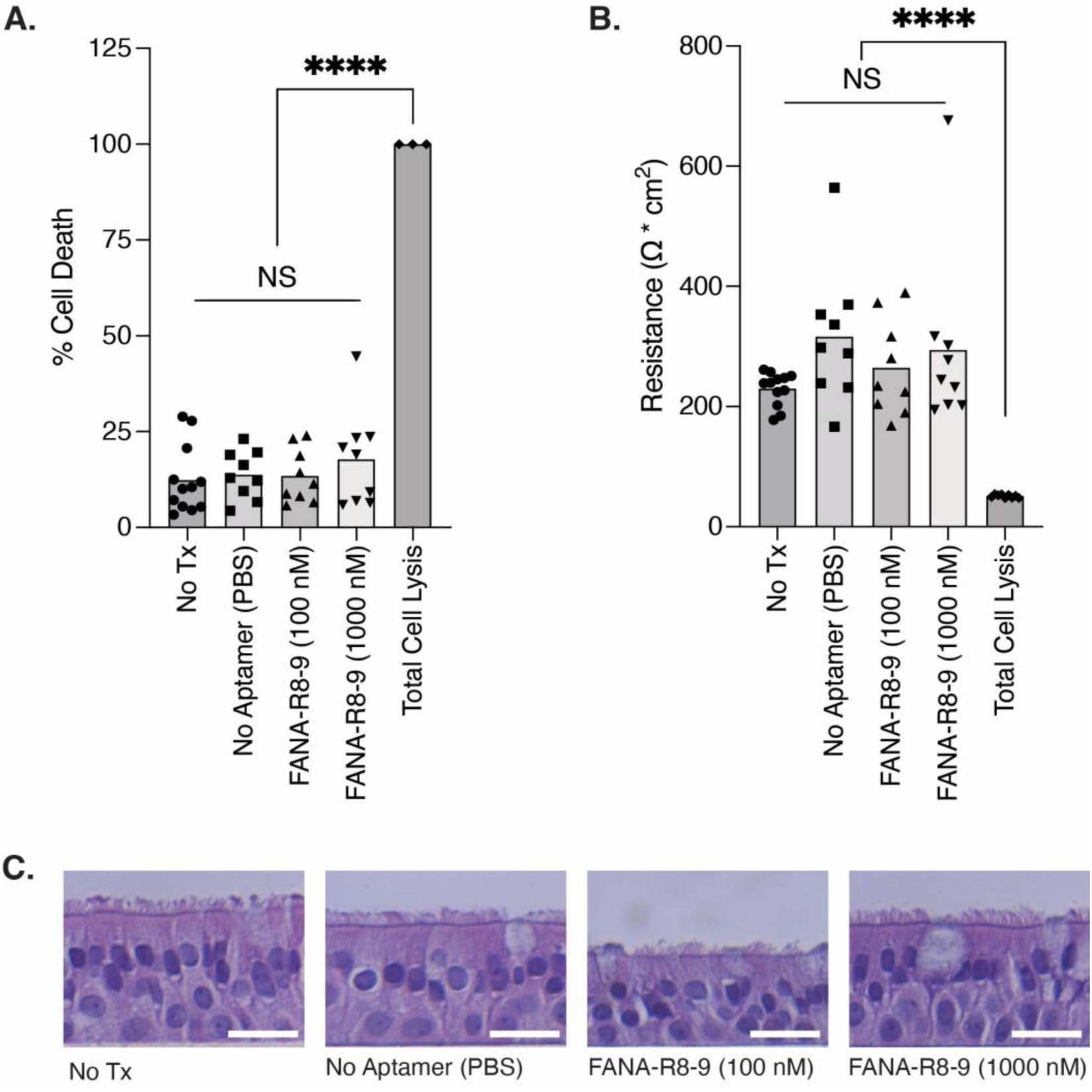
A lack of overt cytopathic effects in HAE cultures following FANA-R8-9 aptamer treatment. To quantify aptamer-induced cytopathic effects, HAE cultures were first inoculated with the indicated treatments at 0, 24, and 48 hrs, (as outlined in Figure 2A). Lactate dehydrogenase (LDH) released into the apical compartment was then quantified at 72 hours and cell death was calculated as a percent of the average total cell lysis control (**A**). Transepithelial electrical resistance (TEER) was then determined in the same cultures (**B**) prior to fixation. Individual biological replicates from two experiments are shown in A and B (n=3-4 replicates per condition, per donor; 3 donors) with the bar representing the mean. Experimental results were analyzed using one-way ANOVA and data are non-significant (NS) or significant (**** p < 0.0001) where indicated. **C**) Representative histological cross-sections of HAE cultures fixed after TEER measurement for one donor. H&E counterstain. Scale bar = 20 microns.

## Discussion

Since the emergence of SARS-CoV-2, a variety of therapeutics have been utilized to inhibit viral replication and mitigate disease severity including immune modulators, monoclonal antibodies, and antiviral drugs such as remdesivir, molnupiravir, and nirmatrelvir with ritonavir. Nonetheless, developing novel strategies to block SARS-CoV-2, as well as other viral infections for which targeted antivirals do not exist, is paramount. Here, we have focused on aptamer technology, which has proven useful in diagnostics [34–36] and demonstrated clinical success with the approval of pegaptinib (Macugen) for treating age-related macular degeneration in 2004. Aptamers specific for viral targets (*e.g.*, polymerases, integrases, helicases) have also been selected, with some demonstraing antiviral activity (Reviewed in [4,5]). However, aptamers do not efficiently enter cells. Therefore, while aptamer uptake into cells can be improved through interaction with cell surface proteins [37–39] or by coupling to cell-penetrating peptides [40,41], aptamers that target viral surface proteins are likely more feasible as antivirals.

Recently, several aptamers to the spike protein of SARS-CoV-2 have been characterized [6–16; Reviewed in: 17,18]. The majority are composed of DNA with the exceptions being a coupled peptide/DNA aptamer [42], an RNA/2’-fluoro RNA (F-RNA) mixed aptamer [9], and the FANA R8-9 aptamer used in this research [15]. The XNA composition of the latter two aptamers could confer some advantages over natural nucleic acid aptamers (see Introduction), although this remains to be determined. Notably, our pseudovirus infections in Vero cells indicated that FANA controls (Fixed FANA and FANA-IN-1.1) were more inhibitory than DNA (Fixed DNA), but less inhibitory than FANA-R8-9. However, none of these controls exhibited stable binding to pseudovirus material, or a significant effect on SARS-CoV-2 infection in our HAE assays. One explanation is that the “weak” binding of FANA to the pseudovirus, that cannot be measured in the assay, is enough to interfere with entry, thus leading to a lower signal. A second possibility is that FANA interferes with replication or transcription of the VSV-derived virus. Remarkably, while studying the ability of HIV genome-targeting FANA antisense oligonucleotides (ASOs) to block HIV infection, researchers not only observed sequence-specific inhibition of HIV infection, but also a non-specific inhibitory component [32]. Furthermore, the inhibitory nature of catabolic metabolites of FANA oligonucleotide, including 2’-deoxy-2’-fluoroarabinonucleosides, against some viruses has been reported [43–45]. Consistent with these previous reports, we have also observed sequence-independent inhibition of HIV by FANA nucleic acids similar in size to FANA-R8-9 and the controls used in the current experiments. Although the cause of the observed non-specific inhibition of pseudovirus infections in Vero cells remains unclear, the potential useful antiviral effects of FANA and other XNAs warrant further analysis.

Importantly, beyond potential XNA-mediated effects, SARS-CoV-2 spike aptamers have been shown to block pseudovirus or live virus [9] infections in cell culture and the respiratory tract of ACE2 transgenic mice [16]. However, there is a clear need to examine aptamers in models that more closely mimic human physiology, an approach already being used to test other potential SARS-CoV-2 inhibitors [46]. Here, we have extended our previous work and demonstrated FANA-R8-9 is not only capable of blocking spike-bearing pseudovirus particles, but also SARS-CoV-2 infection in a physiologically-relevant model of human airway epithelium. FANA-R8-9 was able to reduce infection when mixed with virus prior to inoculation and several hours after viral infection. To the extent that the HAE model mimics the human airway, this suggests that administering aptamer after infection could prevent virus spread and highlights the potential for using aptamers to treat COVID-19 and other respiratory infections. Notably, we did not observe any cytopathic effects, gross morphological changes, or loss of epithelial barrier integrity after 72 hrs of aptamer treatment. While these data are not surprising since we do not expect FANA-R8-9 to be readily taken up by the epithelial cells in the HAE model, the impacts of FANA or FANA-R8-9 on the mucus barrier and underlying cells remain to be fully characterized.

SARS-CoV-2 spike, like other viral glycoproteins, is a primary target of the humoral immune response which can drive glycoprotein sequence evolution over time. As a result, aptamers selected against the glycoprotein (spike) of one virus strain may not inhibit others. Here, FANA-R8-9 was initially selected for binding to the Wuhan virus RBD; however, our data indicate that FANA-R8-9 also effectively inhibits the Delta variant. Indeed, FANA-R8-9 has significantly stronger binding affinity to Delta RBD (23.5 ± 4.6 nM vs. 1.4 ± 0.4 nM, for Wuhan and Delta, respectively) [14]. This enhanced binding may explain why FANA-R8-9 was even more potent against Delta in the HAE assays, although FANA-R8-9 significantly inhibited both strains. Notably, the RBD domains of Wuhan and Delta differ by only 2 amino acids (L452R andT478K) with the changes in Delta resulting in two additional positive charges in the RBD. These additional charges may help enhance the binding of nucleic acid which is strongly negatively charged. In contrast to Delta, Omicron variants typically have a dozen or more amino acid changes in the RBD compared to Wuhan and FANA-R8-9 binds much less strongly to Omicron RBD. We are currently working on producing FANA aptamers to Omicron RBD and S1 domains.

## Conclusions

Overall, nucleic acids as therapeutics have been under rapid development with several FDA-approved drugs and more in the pipeline (Reviewed in [47–49]). Most have been small interfering RNAs (siRNAs), ASOs, splice switching RNAs, or mRNAs. To date, siRNA and ASOs directed to respiratory targets have shown some efficacy in animal models. These successes, in conjunction with our current data, promise further development of FANA-R8-9 and other virus-specific aptamers as therapeutics in the near future.

## Declarations

### Availability of data and materials

All data generated or analyzed during this study are included in this published article.

### Authors’ Contributions

NR generated aptamers, performed experiments (pseudovirus particle assays; *en face* staining and imaging of SARS-CoV-2 infection in HAE cultures; cytotoxicity assays in HAE cultures), analyzed and curated data. WL performed experiments with SARS-CoV-2 virus. MAI performed experiments (*en face* staining and imaging of SARS-CoV-2 infection in HAE cultures; imaging of H&E stained sections) and analyzed and curated data. JML prepared apical HAE culture secretions for use in degradation experiments. EIA prepared pseudovirus particles and established methods for pseudovirus particle assays. XZ supervised all ABSL3 work and acquired funding. MAS conceptualized the study, designed methods, supervised experiments utilizing HAE cultures, acquired funding, and wrote the manuscript. JJD conceptualized the study, designed methods, supervised the generation of aptamers and experiments utilizing pseudovirus particles, performed experiments (aptamer binding to pseudovirus particles; aptamer degradation assays), analyzed and curated data, acquired funding, and wrote the manuscript. All authors read and approved the final manuscript.

## Acknowledgments

We thank Philipp Holliger (Cambridge Biomedical Campus, Cambridge, UK) for the D4K enzyme used to make FANA. We are also grateful to the directors, managers, and staff of the New York University Experimental Pathology Research Laboratory. The following reagents were obtained through BEI Resources, NIAID, NIH: African Green Monkey Kidney Epithelial Cells (Vero E6) expressing high endogenous angiotensin-converting enzyme 2 (NR-53726); SARS-Related Coronavirus 2, isolate New York-PV08410/2020 (NR-53514); and SARS-Related Coronavirus 2, isolate hCoV-19/USA/PHC658/2021 (lineage B.1.617.2; Delta Variant; NR-55611), contributed by Drs. Richard Webby and Anami Patel.

